# Discovery of perturbation gene targets via free text metadata mining in Gene Expression Omnibus

**DOI:** 10.1101/150896

**Authors:** Djordje Djordjevic, Joshua Y. S. Tang, Yun Xin Chen, Shu Lun Shannon Kwan, Raymond W. K. Ling, Gordon Qian, Chelsea Y. Y. Woo, Samuel J. Ellis, Joshua W. K. Ho

## Abstract

There exists over 2.5 million publicly available gene expression samples across 101,000 data series in NCBI’s Gene Expression Omnibus (GEO) database. Due to the lack of the use of standardised ontology terms in GEO’s free text metadata to annotate the experimental type and sample type, this database remains difficult to harness computationally without significant manual intervention.

In this work, we present an interactive R/Shiny tool called GEOracle that utilises text mining and machine learning techniques to automatically identify perturbation experiments, group treatment and control samples and perform differential expression. We present applications of GEOracle to discover conserved signalling pathway target genes and identify an organ specific gene regulatory network.

GEOracle is effective in discovering perturbation gene targets in GEO by harnessing its free text metadata. Its effectiveness and applicability has been demonstrated by cross validation and two real-life case studies. It opens up new avenues to unlock the gene regulatory information embedded inside large biological databases such as GEO. GEOracle is available at https://github.com/VCCRI/GEOracle.

## 1. Introduction

The NCBI Gene Expression Omnibus (GEO) is one of the largest public repositories for genome-wide omic data, including mostly transcriptomic data [1]. As of August 2018, GEO contains over 101,136 data series (GSE), consisting of over 2.5 million individual gene expression samples (GSM). This database harbours biological insights that are not apparent when studying each data set individually [11]. Several packages are available to programmatically access GEO data, including GEOquery [3], GEOmetadb [14], compendiumdb [7] and shinyGEO [5], allowing keyword based search and download of GSE and GSM, with few standard analysis options.

One major challenge in effectively reusing public gene expression data is the availability of good quality metadata. The need for standardisation of metadata is the reason for the development of the Minimum Information About a Microarray Experiment (MIAME) standard [2], and more recently the MINSEQE standards for sequencing data [11]. While some fields in GEO metadata use controlled vocabularies (*e.g.*, species name, gene symbols), the majority of the metadata appears as free text, describing the context of samples (*e.g.*, tissue type or developmental stage) and the experimental design (*e.g.*, perturbation experiment). Although this free text is often readily interpretable by humans, there is no simple means to process this information from GEO in an automated fashion. Ultimately this imposes a major limitation on effectively re-using the huge amount of public data in GEO [11]. While we believe it is important to push for the use of standard annotations, we nonetheless wish to reuse the large amount of data that exists in GEO.

A gene expression experiment can typically be classified based on its experimental design (*e.g.*, perturbation, time-series and case-control experiments). In many cases, data sets from perturbation experiments (*e.g.*, gene knock-out, signalling stimulation, or physical stimulation) are valuable because they allow us to identify the set of genes that are causally downstream of the perturbation agent. This has important applications in determining signalling pathway targets and regulatory networks [8, 4, 13, 12]. There are tens of thousands of perturbation studies in GEO, likely containing millions of experimentally determined perturbation data. Nonetheless, currently there is no simple way to determine whether a GSE contains perturbation data. Furthermore, even when a GSE is known to contain perturbation data, it is not trivial to automatically match the treatment samples with their respective control samples since a single GSE may contains multiple treatment and control groups.

In light of these challenges, we propose to harness the wealth of information embedded in the free text metadata associated with each GEO entry. In GEO, free text metadata include GSE title, GSE experiment type, GSE summary, GSE overall design, GSM title, GSM characteristics, and GSM treatment protocol. These metadata fields are considered ‘free text’ because submitters are not required or do not customarily use standardised ontology terms. In GEO, only very few fields enforce the use of standard ontology terms, such as gene symbols, and organism. Our approach is to use text mining and machine learning techniques to classify GSE that contain perturbation data, and to identify and match the treatment and control samples (GSM) in a perturbation data set. Text mining of free text metadata has previously been used to identify related experiments through semantic similarity [6], and to automatically process large amounts of the GEO database with limited quality control or user oversight [15].

Using our R Shiny tool called GEOracle, we can quickly annotate many perturbation experiments from GEO in a semi-automated fashion with full user control. GEOracle then performs differential expression analysis to identify gene targets of the perturbation agent.

## 2. Methods

The GEOracle workflow mimics the steps a bioinformatician would employ when identifying and analysing perturbation data in GEO. GEOracle uses GEOmetaDB [14] to access GEO metadata. Given a list of GSE accession number (or results from a GEO keyword search), GEOracle will carry out the following six steps:

1. Classification of perturbation GSE
2. Grouping of GSM into replicate sample groups
3. Classification of each GSM sample group as ‘perturbation’ or ‘control’
4. Matching each ‘perturbation’ sample group with the most relevant ‘control’ group
5. Manual inspection and adjustment of predicted annotation through the GEOracle graphical user interface
6. Automated differential gene expression analysis

In the rest of this section we describe the methodology for performing these steps and evaluate GEOracle’s performance on manually curated training and test sets.

### 2.1. Step 1: Classification of perturbation GSE data sets

To build a classifier for identifying perturbation experiments, we curated a training set of 277 randomly selected GSE IDs, which we manually annotated with the experimental design (Additional File 1). Based on 31 manually defined textual features from the free text metadata that can differentiate perturbation experiments (including keywords such as ‘knockout’, ‘KO’, ‘wildtype’, ‘WT’, ‘null’, ‘-/-’, ‘transgenic’, ‘‘TG’; Additional File 2), a support vector machine (SVM) classifier was built to predict perturbation GSE. When evaluated on our training set by 100 rounds of 10-fold cross-validation with internal feature selection, we found that the use of a SVM with a radial basis function (RBF) kernel produced the highest mean Area Under the Receiver Operating Characteristic curve (AUROC) of 0.89, suggesting high sensitivity and specificity (Figure 1)

**Figure 1:**
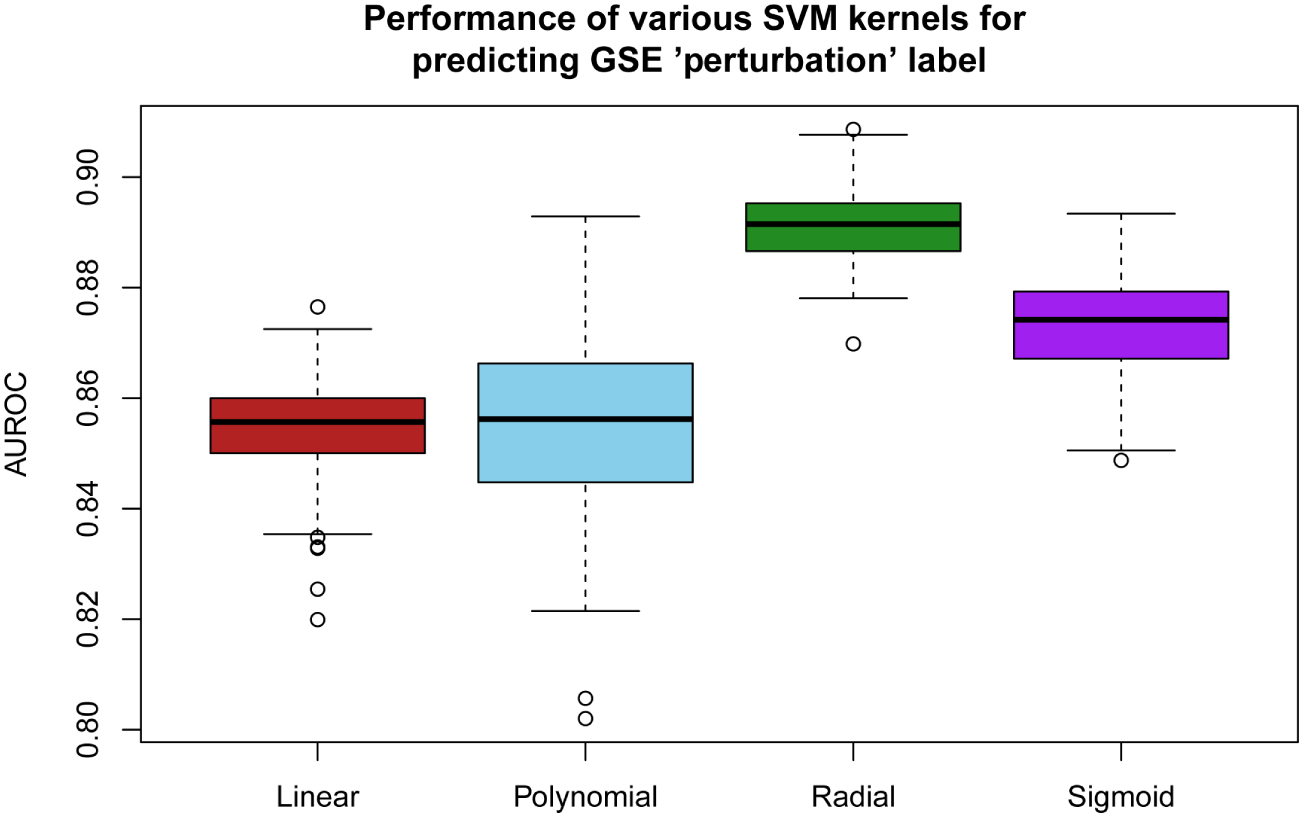
Comparison of the performance of different SVM kernels for predicting GSE ‘Perturbation’ label based on the manually curated training set of 277 GSE IDs. Shown are boxplots of the Area Under the Receiver Operating Characteristic (AUROC) curve from 100 repetitions of 10-fold cross-validation. The radial basis function kernel has the best performance.

### 2.2. Step 2: Grouping of replicate GSM samples

Once a GSE is identified as a perturbation GSE, GEOracle will then automatically group samples (GSM) into replicates. We manually curated a second set of 73 perturbation GSE. We annotated the 832 constituent GSM samples into 259 groups labelled as ‘perturbation’ or ‘control’, and paired the ‘perturbation’ sample groups with their appropriate ‘control’ groups (Additional File 3).

For each identified perturbation GSE, GEOracle groups replicate samples using the available GSM metadata. Replicates could mean biological or technical replicates that together form a unit of analysis for differential expression. GSM titles are processed via a series of string manipulations to remove replicate identifiers and tokenise the titles. A simple hierarchical clustering approach is used, based on Gower distance between tokenised GSM titles, with the tree cut at height 0, resulting in identical GSM titles being assigned to one cluster. The same approach is applied to GSM characteristics to produce a second clustering of samples. Based on these two sample clusterings, we identify the most valid clustering outcome and assign confidences to the output, removing datasets with insufficient metadata or invalid clustering results from further analysis (Figure 2a).

**Figure 2:**
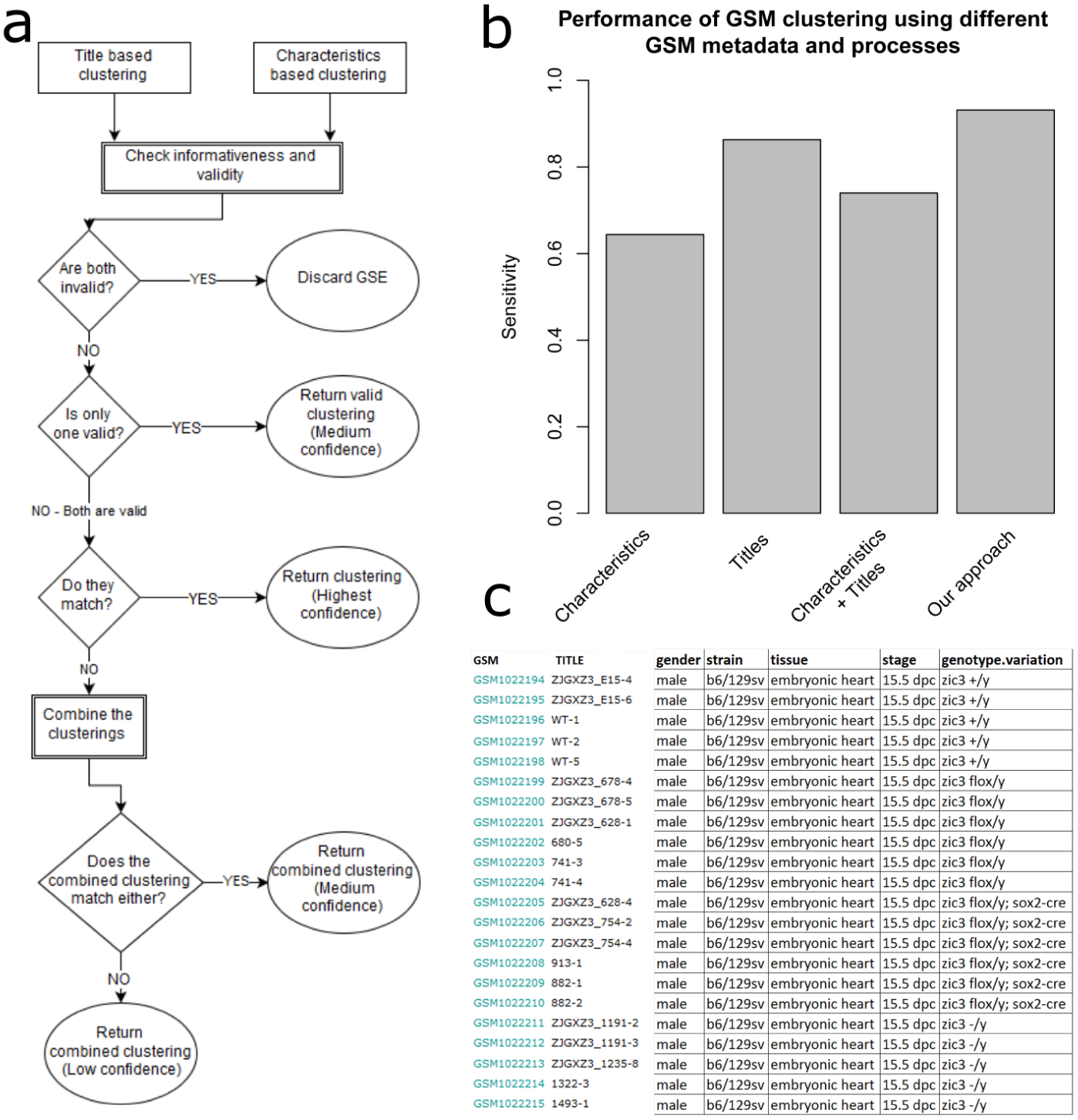
Grouping of GSM samples into groups of replicates. (a) The logic flow for assessing the most valid clustering of GSM samples. This schematic diagram shows the decision making process during the multi-stage clustering procedure that combines information from the GSM titles and characteristics. Informativeness and validity means that there is more than 1 cluster in the GSE and that there are fewer than N clusters, where N is the number of GSM in the GSE. (b) Comparing the performance of clustering using GSM titles and characteristics. Shown is the relative sensitivity of different clustering methods, using GSM characteristics only, GSM titles only, a simple concatenation of GSM characteristics and titles and our multi-stage clustering approach. (c) Sample titles from GSE41674. Title based clustering was not able to correctly cluster this GSE, whereas GEOracle’s multi-stage clustering approach could, by utilizing the information in the GSM characteristics.

Our multi-stage clustering approach produces a grouping sensitivity of 93.2% at the GSE level (meaning every sample in a GSE must be correctly grouped for that GSE to be considered a positive result) based on our training set. All incorrectly clustered GSE can be explained by typographical errors and other anomalies in the metadata. This was an improvement over more naive clustering approaches, based solely on the GSM characteristics, GSM titles, or a simple concatenation of the two, producing sensitivities of 64.4%, 86.3% and 74% respectively (Figure 2b). Although samples can often be grouped by either the titles or the characteristics, the process of deciding which information to use is non-trivial. Figure 2c shows a complex example where a simple concatenation of GSM titles with GSM characteristics fails to group samples correctly, while our multi-stage decision process succeeds.

### 2.3. Step 3: Labelling of GSM sample groups

After grouping GSM samples into groups of replicates, GEOracle proceeds to use a SVM to classify whether each of these replicate group belongs to the ‘treatment group’ or the ‘control group’ in a perturbation experiment. Both the GSM titles and characteristics were analysed for the presence of 33 textual features that represent molecular concepts that can differentiate ‘perturbation’ from ‘control’ samples (Additional File 4). Using cross-validation, we found the linear kernel for the SVM gave the best results, with a prediction sensitivity of 73.3% (Figure 3). This is what we called the raw prediction.

**Figure 3:**
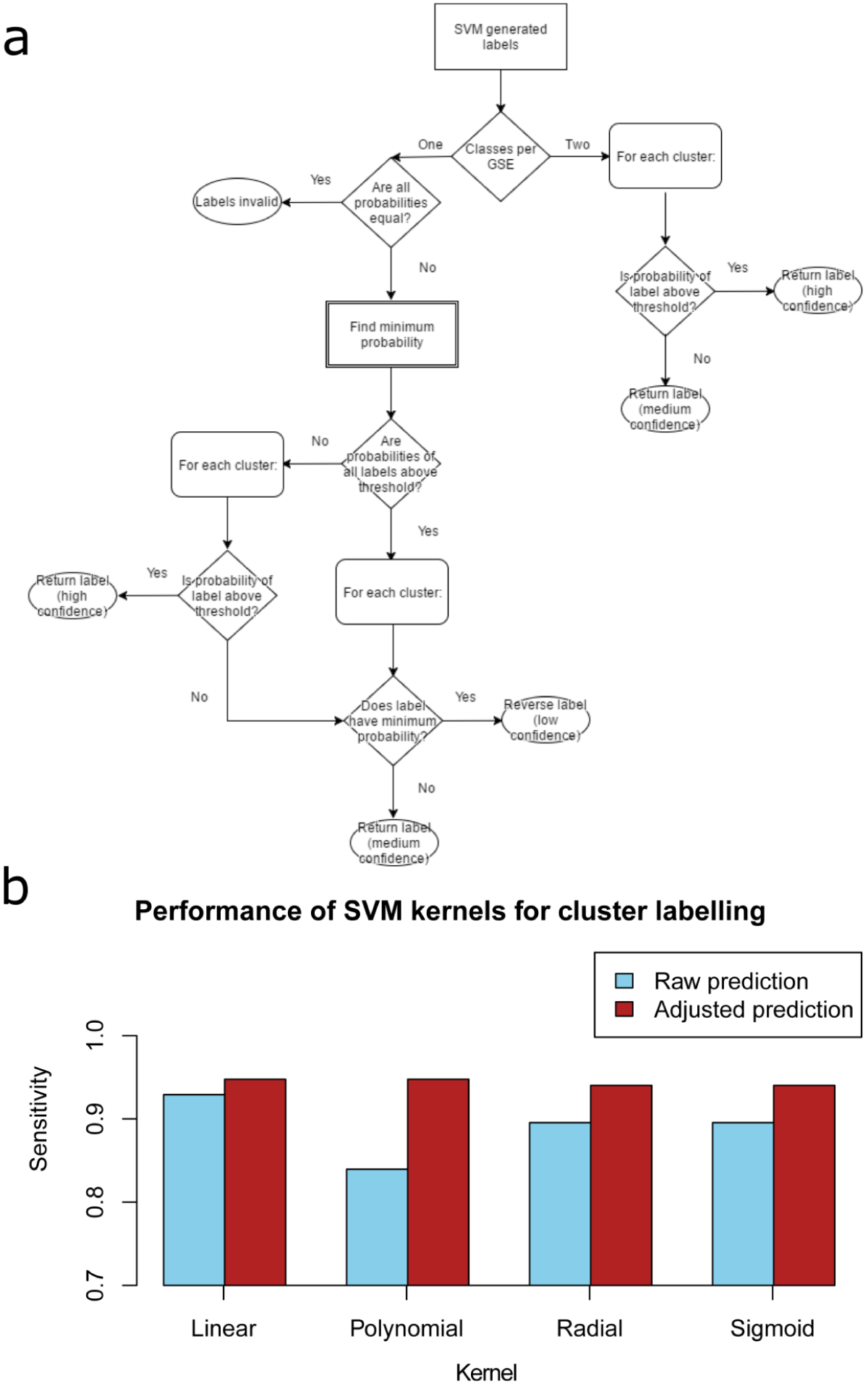
Labelling of GSM. (a) The logic flow for assessing the most valid label for a cluster of GSM. This schematic diagram shows the decision making process for fixing labels (‘perturbation’ or ‘control’) predicted by the SVM based on textual features. This process is particularly important when only one cluster label is generated for every cluster in a GSE. (b) Comparing the performance of different SVM kernels to predict the label of GSM clusters (‘perturbation’ vs ‘control’). Sensitivity is calculated as the fraction of GSE for which the GEOracle output perfectly matches the manually annotated set of 73 GSE. Shown is sensitivity calculated on the raw label predictions (blue) and after cluster label adjustment (red).

To further increase the classification performance, we adjust the predicted labels of some groups when only one label is predicted for all samples in a GSE. A confidence associated with the final outcome of group labelling is determined (Figure 3a). After this adjustment, we can achieve a sensitivity of 94.6% for group classification at the GSE level. This is a large improvement over the 73.3% sensitivity produced by the basic approach of choosing the highest scoring label based on the occurrence of the subset of 20 features that unambiguously distinguish between ‘perturbation’ and ‘control’ samples (Figure 3b).

### 2.4. Step 4: Matching perturbation with control groups

Next, GEOracle matches each perturbation group to the most appropriate control group. GEOracle matches each predicted ‘perturbation’ group to the ‘control’ group with the lowest Gower distance based on the tokenised GSM titles and characteristics, and determines the confidence for each pairing of groups (Figure 4). We observe a sensitivity of 83.1% for group matching at the GSE level. Furthermore, we attempt to determine the identity of the perturbation agent and perturbation direction for each group pair by searching for gene names and keywords in the GSM and GSE metadata. The keywords used represent the concepts of addition (*i.e.*, ‘overexpress’) and removal (*i.e.*, ‘knockout’) of a perturbation agent. The direction with the most keyword matches becomes the assigned direction.

**Figure 4:**
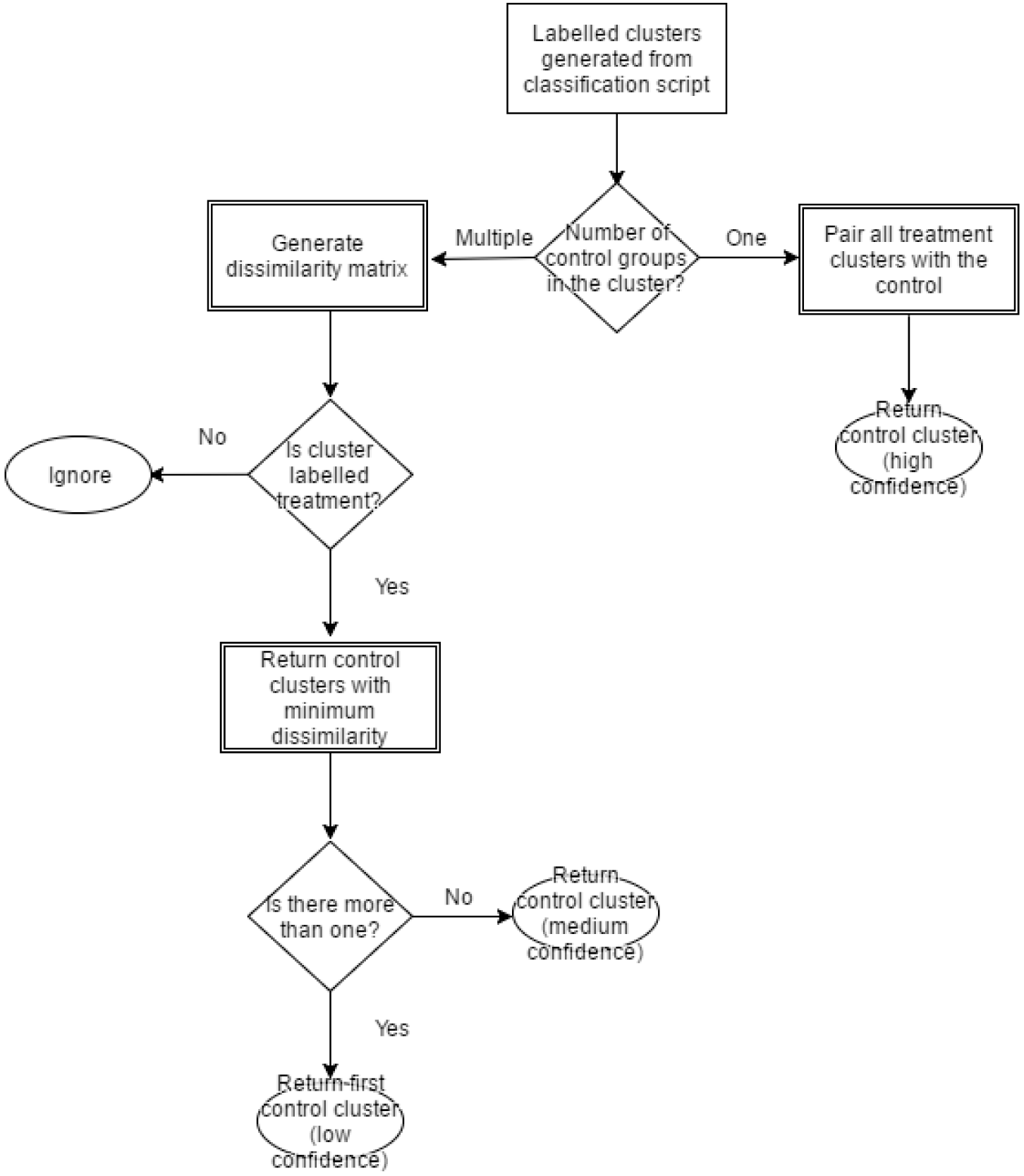
The logic flow for pairing labelled clusters. This schematic diagram shows the decision making process for matching a ‘perturbation’ cluster which its closest ‘control’ cluster. This can be non-trivial when multiple ‘control’ clusters exist within a GSE. Dissimilarity is Gower distance.

### 2.5. Step 5: Manual adjustment using the GEOracle graphical user interface

The GEOracle interface (Figure 5) guides users through the entire process. Importantly the interface allows a user to manually verify and adjust all the details of the predicted GSM labels and pairings, and create their own pairings from all GSM within each GSE. This allows the user to be 100% confident in the setup of samples for differential expression analysis.

**Figure 5:**
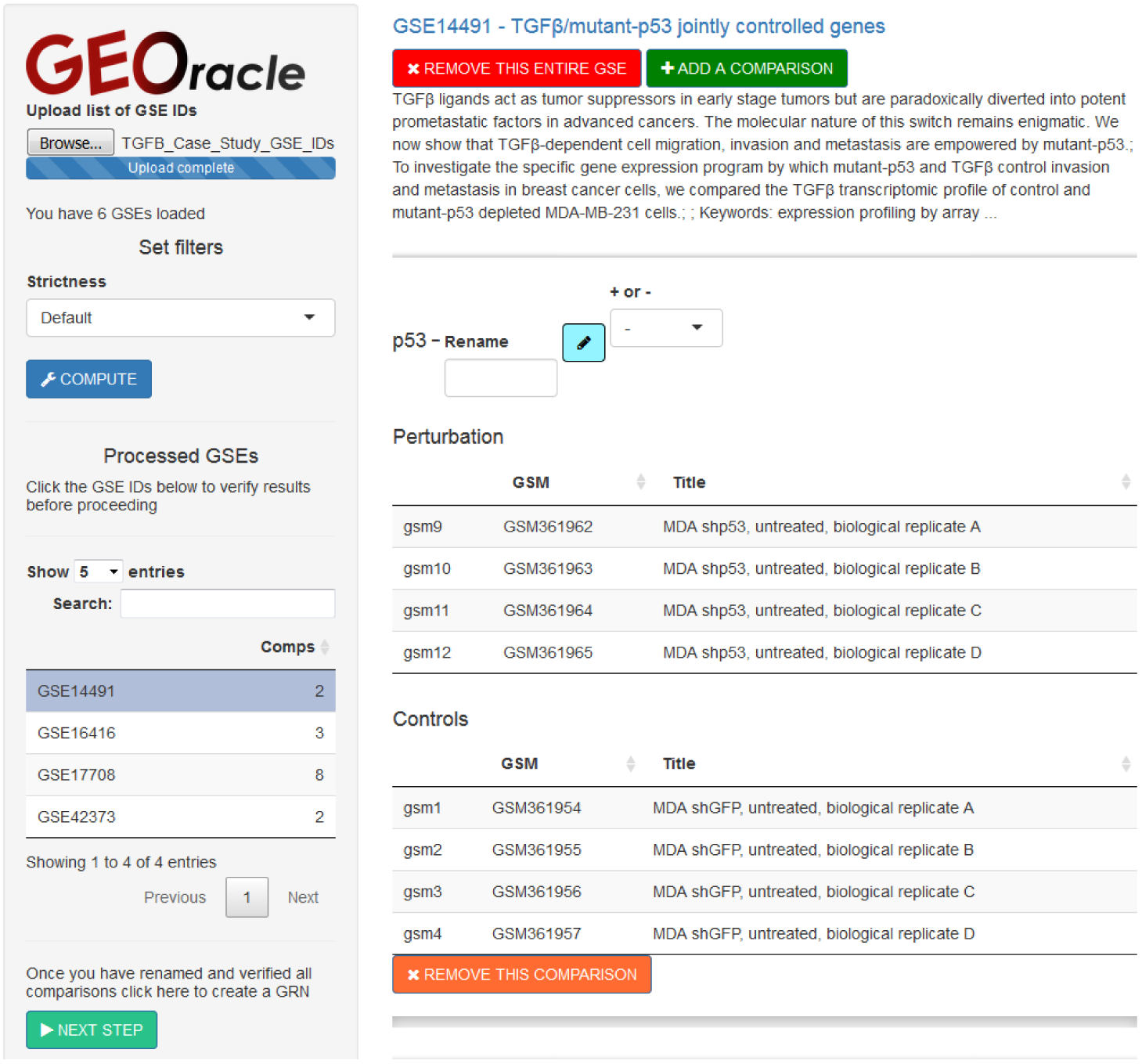
The GEOracle user interface.

### 2.6. Step 6: Automated differential expression analysis

The paired ‘perturbation’ and ‘control’ groups are then used to compute differential gene expression using GEO2R, which implements the limma pipeline [10]. The results can then be downloaded by the user. GEOracle is currently tailored for microarray data analysis as this is the most prevalent data type in GEO, but it can be extended to analyse RNA-seq data or even other functional genomic data sets such as ChIP-seq.

## 3. Case studies

### A conserved response to TGFβ stimulation in human cells

We used GEOracle to process six GSE containing TGF*β* perturbation experiments (Additional File 5) and discovered the consensus target genes of TGF*β* signalling stimulation in human cells (Additional File 6 describes the process in detail). The total time required for classifying the GSE and GSM groups, matching the treatment and control samples, manually verifying the results, downloading the gene expression data from GEO and performing differential expression analysis is less than 12 minutes. This analysis required minimal human intervention and essentially no bioinformatics expertise.

Based on these results we could identify a consensus TGF*β* target gene signature in human cells consisting of 82 genes (Figure 6). Many of the observed transcriptional changes matched the literature about the TGF*β* pathway, including increased transcription of *CTGF*, *JUN*, *JUNB* and *WNT5B*, and repression of *TGFBR3*, *FZD7* and *SPRY1* (Additional File 7). A gene ontology analysis of the 82 genes from the consensus signature using g:Profiler [9] showed significant enrichment for the term ‘response to transforming growth factor beta’, with Benjamini-Hochberg (BH) adjusted p-value of 8.93 × 10*^−^*^08^.

**Figure 6:**
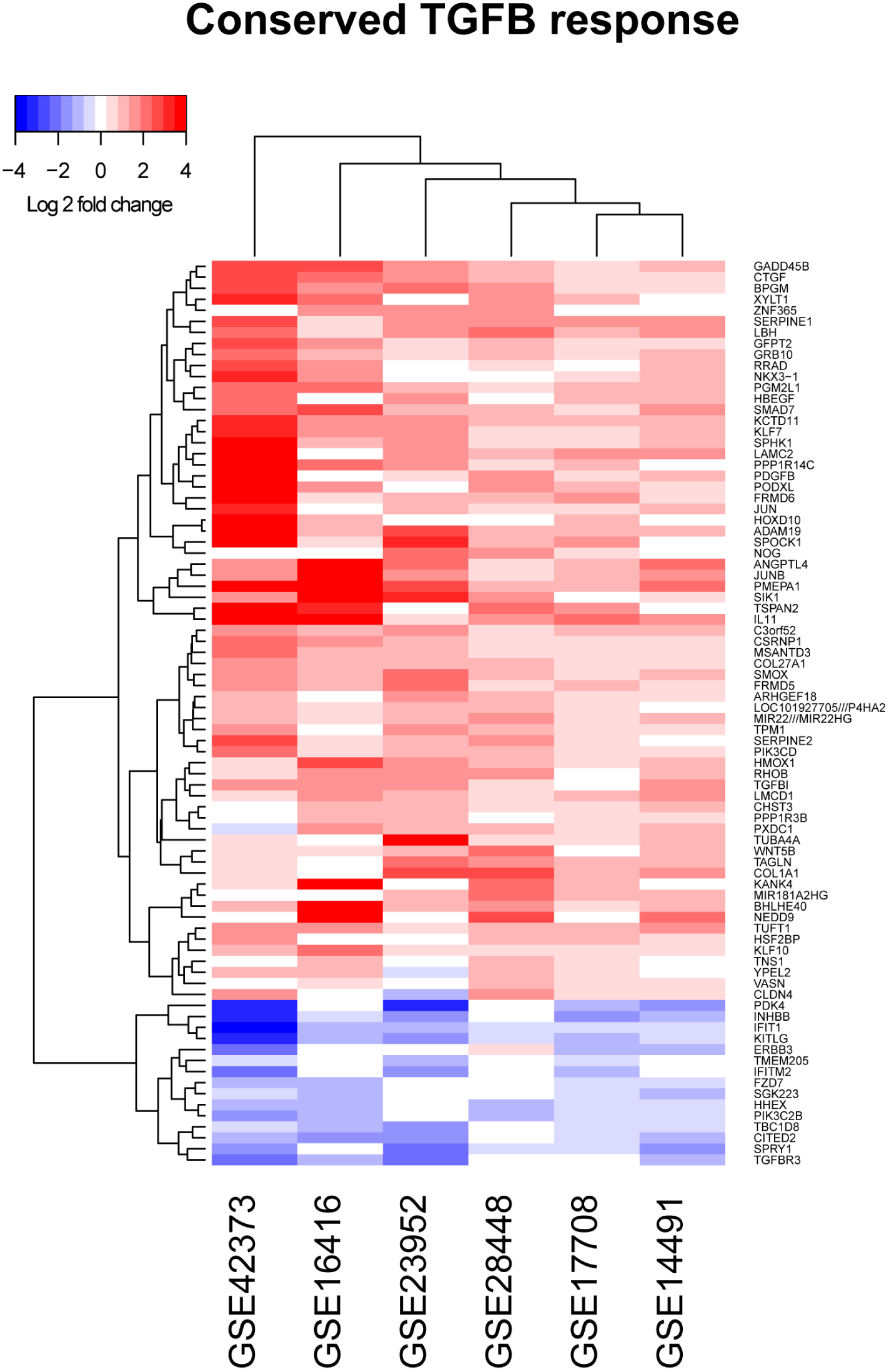
A conserved gene expression signature upon TGF*β* stimulation in human cells. This heat map summarises the genes that are up-regulated (red) and down-regulated (blue) based on various gene expression data sets (GSE).

### 3.1. Identifying a causal gene regulatory network of mouse heart

We further used GEOracle to analyse all perturbation microarray data from mouse cardiac tissues. We searched GEO using the following query: “mus musculus”[Organism] AND (“heart”[MeSH Terms] OR heart[All Fields] OR cardiac[All Fields]) AND (“gse”[Filter] AND “Expression profiling by array”[Filter]). This resulted in 851 GSE (Additional File 8).

Processing these 851 GSE though GEOracle, including manually verifying and modifying the predicted sample comparisons, required approximately 8 hours of user time, again with minimal bioinformatics expertise required. 164 relevant GSE were included for further processing. The primary reason for exclusion was that the tissue profiled was not of cardiac origin or relevance. We obtained significantly differentially expressed genes for 87 genetic perturbations (*i.e.*, gene knockdown or over expression) and 10 non-genetic factors (diet, chemicals etc.) using standard thresholds (absolute log 2 fold change > 1 and BH adjusted P value < 0.05). GEOracle automatically outputs significant differentially expressed genes as an edge list for gene regulatory network construction. From the genetic perturbation experiments we constructed a gene regulatory network of 23,347 causal and directed relationships between 9,152 genes (Figure 7). Of these 14,120 were activating relationships and 9,681 were inhibitory. This case study illustrates how we can construct a large organ-specific gene regulatory network from published experimental perturbation data in GEO.

**Figure 7:**
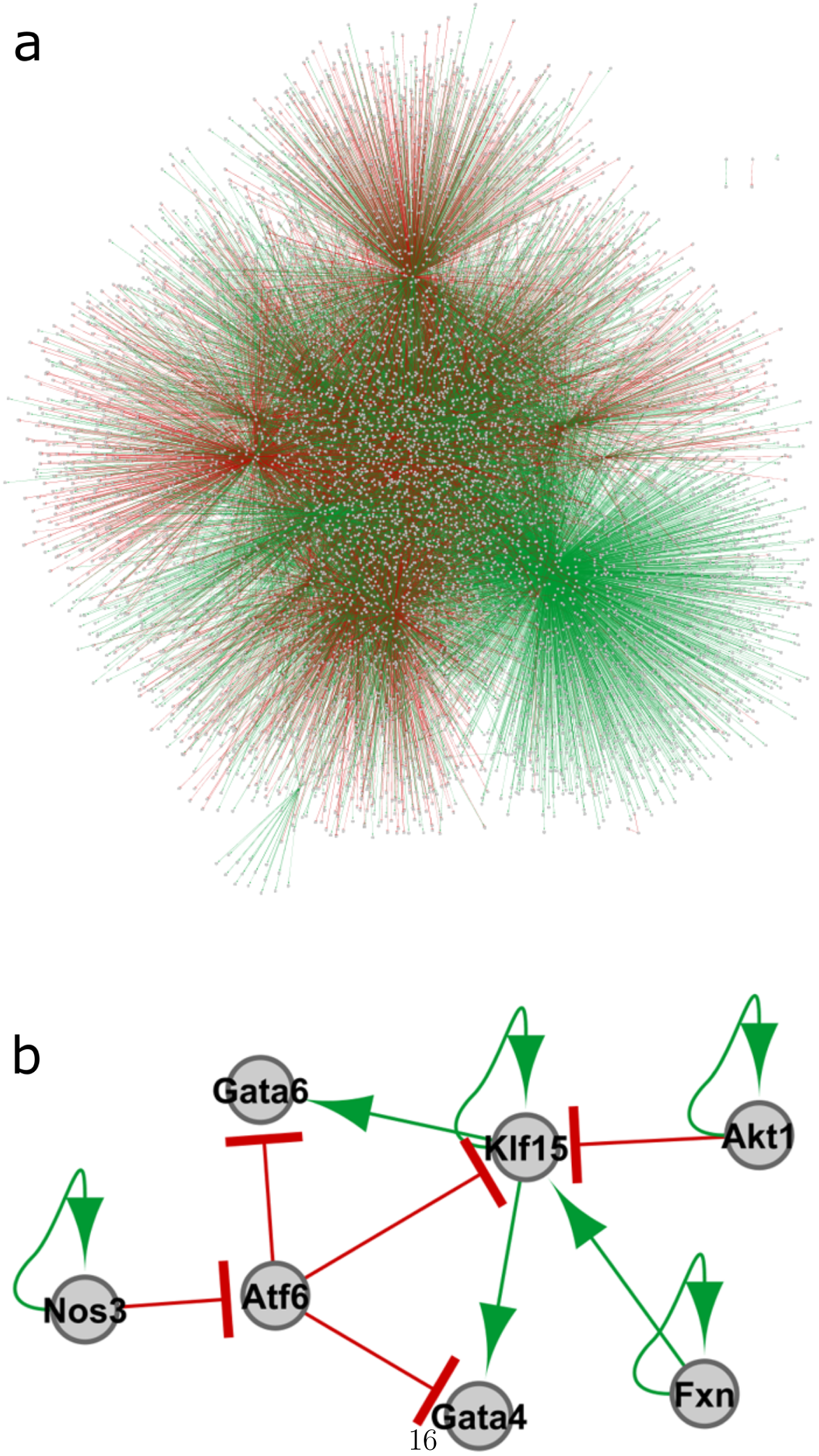
A mouse heart gene regulatory network inferred based on perturbation data mined by GEOracle. (a) Using causal gene regulatory evidence extracted from 97 perturbation data sets by GEOracle, we can construct a murine cardiac gene regulatory network of 23,347 causal edges between 9,152 genes. (b) A sub-network of 7 genes focusing the

## 4. Discussion

In this paper, we present a general framework and an open source R/shiny tool for identifying and processing perturbation gene expression data sets through free text metadata mining. Using GEOracle, we can automatically identify GEO data sets that use a perturbation experimental design, and identify the treatment and matching control groups within these data sets. Our cross validation experiments demonstrate that GEOracle achieve high sensitivity and specificity. Furthermore, GEOracle provide a user-friendly Shiny interface to enable users to manually check and confirm the classification made by GEOracle.

One major innovation of this approach is that GEOracle turns free text metadata in GEO into useful structured information that enables automated mining of a large amount of perturbation gene expression experimental data in GEO. As demonstrated by our two case studies, GEOracle can be used effectively to identify downstream target genes from multiple data sets in GEO.

The current version of GEOracle is implemented in as an R package because it is a widely used programming language for gene expression bioinformatics analysis. Even thought the performance is reasonable for up to hundreds of GSE, it will likely not be able to deal with a larger proportion of GSE in GEO. In the future, we plan to scale up GEOracle by utilising cloud-based parallel and distributed computing technology such as Hadoop MapReduce and Spark.

## Acknowledgements

This work was supported in part by funds from the New South Wales Ministry of Health, a National Health and Medical Research Council Career Development Fellowship (1105271 to JWKH), and a National Heart Foundation Future Leader Fellowship (100848 to JWKH). We thank members of the Ho Laboratory for their valuable comments.

